# Higher surfaces of a crop in the landscape increase outbreak risks the following growing season

**DOI:** 10.1101/641555

**Authors:** T. Delaune, R. Ballot, C. Sausse, I. Felix, M. Chen, F. Maupas, M. Valantin-Morison, D. Makowski, C. Barbu

**Affiliations:** UMR Agronomy, INRA, AgroParisTech, Université Paris-Saclay, BP 78850 Thiverval-Grignon, France; Institut Technique de la Betterave, BP 75008 Paris, France; Terres Inovia, Centre de Grignon, BP 78850 Thiverval-Grignon, France; ARVALIS Institut du végétal, Domaine expérimental du Chaumoy, BP 18570 Le Subdray, France

**Author notes:** Corresponding author: Corentin Barbu.

## Abstract

The use of fungicides and insecticides by farmers represents a major threat to biodiversity^1^, endangering agriculture itself^2,3^. Landscapes could be designed^4^ to take advantage of the dependencies of pests^5,6^, pathogens^7^ and their natural enemies^8^ on landscape elements. However, the complexity of the interactions makes it difficult to establish general rules. Despite initial enthusiasm^9^, the many studies opposing cultivated and semi-natural habitats have not revealed a homogeneous response of pests^10^ and pathogens^11^ to semi-natural habitats. In addition, the question of the impact of crop diversity on pests and pathogens remains largely open^12^. Based on about half a million observations over nine years on 30 major field crop pests and pathogens spread over all latitudes of metropolitan France, we show that the outbreak risk increases with the area of the host crop in the landscape the previous growing season. The impact on the risk of the host crop area the ongoing growing season diverges between animal pests and pathogens. We also confirm that woodlands, scrublands, hedgerows and grasslands do not have a consistent effect over the spectrum of pests. The spatial and temporal distribution of the resource, the host crop, generally prevails over the effects of potential alternative habitats. Territorial and temporal coordination generally promoting crop diversity but excluding a crop at risk a given year may prove to be key levers for reducing pesticide use^14^.

---

During the past decade, the growing awareness of environmental hazards associated with agricultural intensification^1,13^ has motivated abundant studies on alternative agronomic levers to alleviate crop damage caused by pests.^14^. The focus of such research has progressively shifted from its historical focus on the field to the landscape^9,11^ with an emphasis on the opposition between crop and non-crop elements^10,15^ to design integrated pest management strategies at the landscape scale^16^. Studies opposing the cropland to semi-natural habitats failed to define a general rule of thumb regarding the regulation of animal pest epidemics^10,17^. Complex trade-offs between the impact of such landscape elements on both the life cycle of animal pests and their natural enemies have been pointed out^8,18,19^. No clear agreement emerges either for the management of crop pathogens as empirical studies are scarce despite repeated calls for landscape level assessments^11,20.^. Here, we assess the impact of the landscape composition on 30 animal pests or pathogens of 6 annual crops (wheat, barley, maize, potato, oilseed rape and sugar beet). We consider separately the host-crop in the landscape the year of the observation and the year before. We also distinguish within semi-natural habitats the woodlands, scrublands, hedgerows and grasslands. For each landscape element we consider four distances at which their influence can be exerted that roughly correspond to potential management units: 200m (the neighbouring field), 1 km (the farm), 5 km (the village) and 10 km (the group of neighbouring villages). Based on observations from the French epidemio-surveillance network (2009-2017), we carried out binomial LASSO generalized linear regressions on the outbreak risk by pest and pathogen as a function of the landscape composition in a total of 39 880 field × year × pest observation series. In these regressions, we controlled for the effects of preceding crops on the observed field, farming system regions and the regional weather conditions. We also test the robustness of our findings comparing alternate model specification.

* * *

The variable selection left only 5 of the 30 considered organisms without perceptible influence from landscape variables. Area of woodlands, scrublands, hedgerows, grasslands, host crop the ongoing growing season, and host crop the previous growing season had a significant impact on the outbreak risk respectively for 19, 9, 10, 13, 15, and 14 organisms (Fig. 1), often in opposite ways, recalling the relevance of considering the impact of landscape composition over a large range of pests and pathogens. We observed coherent results between models with different specifications (SI. 5) at the exception of the one that did control neither for the effect weather conditions : the detection of most landscape effects on pests and pathogens outbreak risk was actually possible only if the weather was accounted for through a year × climatic region factor.

**Figure 1.**
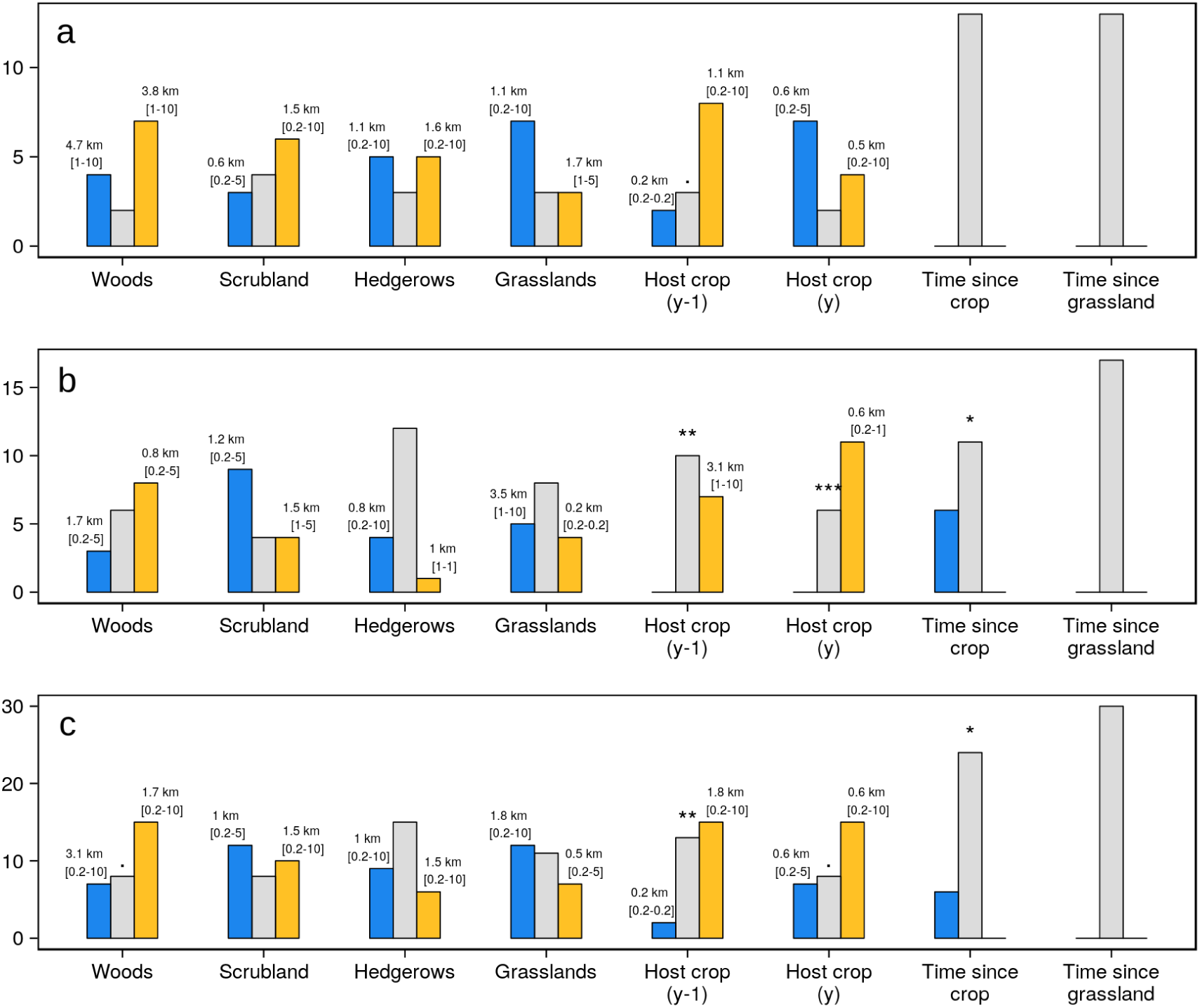
Number of organisms which outbreak risk is correlated positively (orange), negatively (blue), and unaffected (grey) by each of the landscape components for a) pests, b) pathogens, and c) all organisms confounded. Geometric average and range of the most significant distance over the different pests is indicated above the bars. p-value levels for the binomial test comparing the numbers of organisms positively and negatively correlated with an element: “.”: <0.1, “*”: <0.05, “**”: <0.01 and, “***”: <0.001.

To assess the coherence of the landscape composition impact over the pool of organisms studied, we compared the number of organisms positively and negatively impacted by each landscape elements (Fig. 1). The outbreak risk tended to be correlated with the host crop area the previous growing season (Binom. test: P < .01, Fig. 1c) for both pests (Binom. test: P = .055, Fig. 1a) and pathogens (Binom. test: P < .01, Fig. 1b) indicating that for half of the organisms (Fig. 1c), epidemics are likely to occur more frequently in host crop fields if the host crop was largely represented in the surrounding landscape the preceding growing season. Two animal pest outbreak risks were nevertheless negatively correlated with the area of their host crop in the immediate neighborhood the previous growing season evoquing the possibility of occasional regulation by previously attracted natural enemies.

Host crop area the ongoing growing season showed contrasted results between pathogens, generally positively correlated (Binom. test: P < .001, Fig. 1b), and pests, often negatively correlated (Binom. test: P = .27, Fig. 1a), resulting in a statistically significant difference between pests and pathogens regarding the effect direction (Fisher test P < .01). We note that 5 of the 6 tested coleoptera, also univoltine, are negatively correlated with the host crop the ongoing growing season while the 4 tested aphids, notoriously multivoltine and maybe less active in their search of their host crop, are not (SI 6). The average distance at which pathogens are affected by the host crop area was significantly higher (Fisher test P = 0.026) for the previous growing season (3.1 km range across pathogens from 1 to 10 km) than for the ongoing growing season (0.6 km range 0.2 to 1 km, Fig. 1b). Such a difference was not observed for pests, reinforcing the contrast between these groups.

The length of time since the host crop was cultivated on the observed field consistently reduced the outbreak risk for 6 pathogens (Binom. test: P < .05, Fig. 1b). However, no organisms were affected by the time since grassland was cultivated on the field. Overall, the rotation in the field was strikingly less often selected than the area allocated to the host crop in the landscape. The buffer size at which the host crop area mostly affected organisms was largely beyond the field size, averaging 1.1 and 3.1 km for pests and pathogens respectively (Fig. 1a,b). Given the diversity of the current agronomical situations in France, the field level crop rotation seemed to have less impact over the considered pests and pathogens than the landscape over much larger areas than the field. This might not only be due to intrinsic dispersal capacity but also to passive dispersal, notably by machinery or seeds ^21^.

The area of semi-natural elements in the agricultural landscape was very often correlated with the outbreak risk (Fig. 1c). We selected at least one semi-natural elements for most of the organisms (26 over 30) with comparatively more relationships for pests than for pathogens (F. test P < .05). Nevertheless, no trend emerges in the direction of these relationships other than a slight correlation of woods with an increased risk (Binom. test: P<0.1). The data do not support an *a priori* protective effect of the semi-natural habitats against animal pests or diseases.

To grasp the size of the change in epidemiological risk induced by the observed variability in the landscape, we measured the estimated risk change with few (5^th^ percentile) vs. a lot (95^th^ percentile) of each considered element in the landscape (Fig. 2). The area allocated to host crop the previous growing season, already found to impact most organisms (Fig. 1), was associated with the highest average effect size (Fig. 2a). Landscapes with most host crop area the previous growing season are estimated to be on geometric average 8%, range [-2%, +79%], more likely to experience pests or pathogens outbreaks than landscapes with least host crop area the previous growing season. Our estimates of the changes in epidemic risk with landscape compositions are relatively small compared with those induced by contrasting weather conditions as reflected by most risky years on average 53% range [0% – 306 %] more likely to experience outbreaks than least risky years (Fig. 2b).

**Figure 2.**
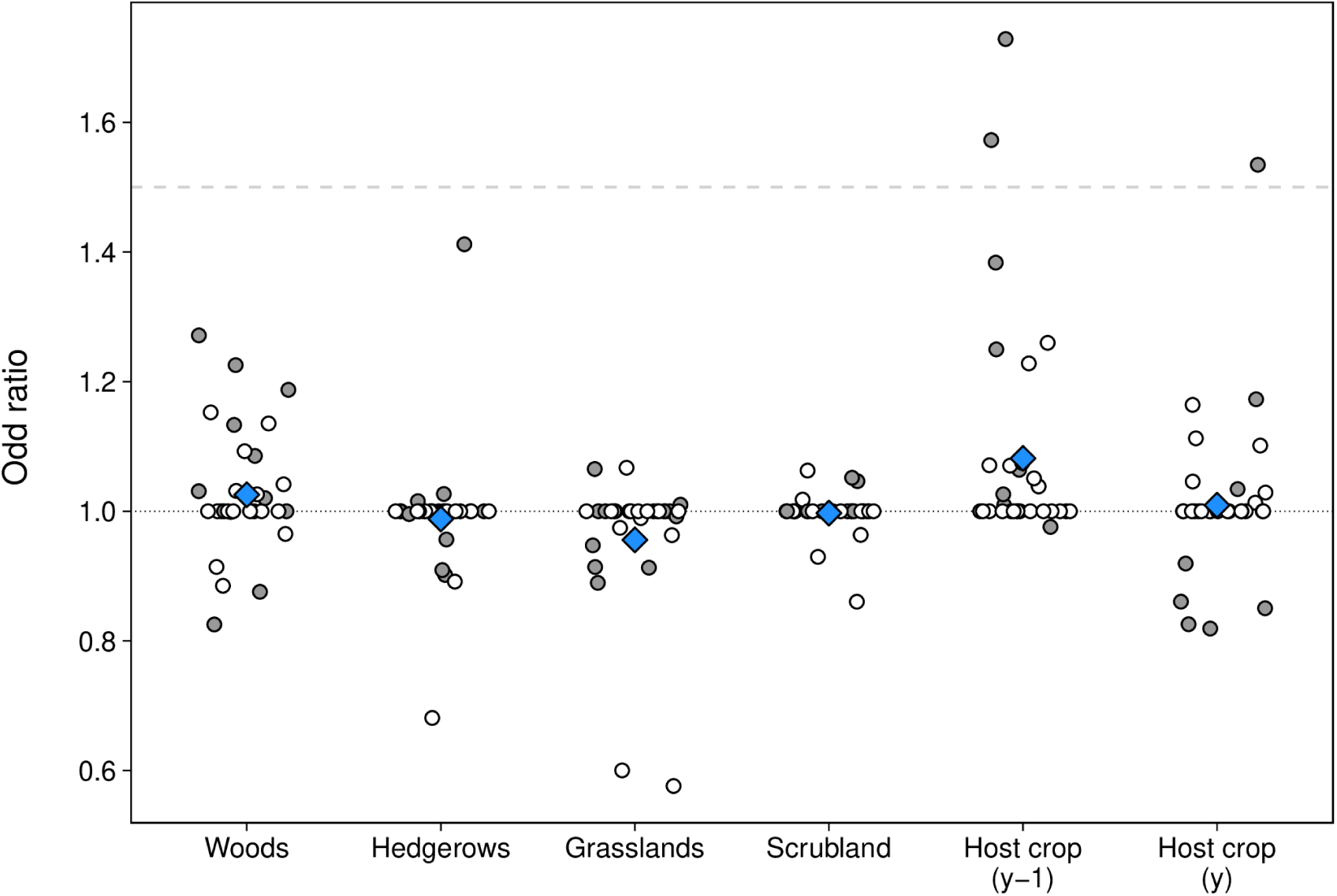
Estimated odd ratios of animal pests (grey dot, n = 13) and pathogens (white dot, n = 17) outbreak risk with each landscape element, taken from the estimate on the full dataset. The odds ratios are taken between the 2.5% and the 97.5% quantile of the observed coverages by the landscape element. Geometric averages of the odd ratios over all organisms are represented by the blue diamonds. The typical effect size of the weather is represented by the dashed grey line: the median over the pests of the year related odds ratio. The year related odds ratio for a pest is the median over the different agro-climatic regions of the odds ratios between the worst and the best year.

* * *

While an accent has been made in the literature on the semi-natural habitats and natural regulations^9,19^, the area allocated to a crop had a much clearer tendency to increase the outbreak risk the following growing season. This suggests that food resources are generally dominant landscape drivers of animal pest but also pathogen population variations. The contrast between animal pests and pathogens in their response to the host crop area the ongoing growing season may be explained by two mechanisms. First, pests unlike pathogens can choose where to land allowing them to cluster on small areas of the host crop^22^. Second, some pests perform only one life cycle per year, preventing epidemic propagation the same year^23,24^.

The semi-natural elements were often correlated or anti-correlated with the outbreak risk but without a consistent trend. This observation already made on insect pests^19^ is here extended to pathogens. Theoretical research on the pathogens show that these areas can serve as barriers but can also present wild hosts facilitating transmission^11,25^. For pests, reviews consistently demonstrated an increased presence of the natural enemies with semi-natural habitats^9,26^ but a reduction of the abundance of pests with semi-natural habitats is less often found^19^. Our results as well as the most recent work^6,10^ suggest that semi-natural spaces can be just as much a needed resource for pests than for their natural enemies.

These tendencies are largely robust to exact specifications of the model but the risk variations induced by the landscape seem on average sizably smaller than those induced by the weather conditions. The estimated effect sizes are nevertheless likely minimums as the data are heterogeneous (quality of the positioning, multiplicity of the experimenters) and the models are simple (neither interactions nor non-linear effects). Lack of statistical power might prevent us to observe some correlations. The generalized use of pesticides in France as in many developed countries^27^ might further mask correlations and even modulate the equilibrium between pests and natural enemies. Despite the cross-validation strategy adopted^28^, the number of tested features increases the risk of selection of spurious correlation or at least errors in the selection of the most correlated spatial scales for any of the 30 individual organisms. As a consequence, we emphasize the interpretation of significant tendencies across organisms (Fig. 2, 3). A species level analysis could benefit from a better consideration of organism functional traits. In particular, one might consider the diversity of host crops of generalists such as *Rhopalosiphum padi*, here observed only on wheat but a pest of most cereals^29^. Finally, more advanced models could consider interactions between landscape elements as well as their configuration^30^.

The trends we identify could be used to reason the protection of the crops at the landscape level. Early warning system tools based on this approach may modulate risk estimates at the plot level and thus limit the systematic use of preventive treatments^31^. As for active landscape design, the management of host plant areas seems both the most influential and the easiest aspect to handle^32^. Beyond increasing the diversity of cultivated crops in the landscape, at least beneficial against pathogens, a rising pest pressure on a crop could be answered by excluding the crop over a large area. This dynamic recommendation of a “blank” year without a particular crop should not be confused with landscape-scale rotations on all crops that could have catastrophic effects on non-pest biodiversity^26^. Organizing at such scales all the stackholders who often mismatch in terms of objectives and perceptions regarding potential benefits of ecosystem services (Kleijn et al., 2019) is a demanding but necessary challenge.

## Material and methods

### Pests and pathogens data

French epidemiological services organise each year the monitoring of a set of fields. Georeferenced, these fields are assigned to a variety of organisations and visited once a week during the cropping season. Several standardized protocols are applied for each surveyed pests of the crop. For each pest we use the results from the protocol with the highest total number of observations (Tab. SI. 2) in the subsystems Vigicultures^®33^ and VIGIBET (ITB – Sugar Beet Research Institute) that cover 17 of the 22 former French administrative regions, covering all latitudes of metropolitan France and approximately two thirds of its territory (Fig. SI.1). These datasets cover the 2009-2017 period. In total, data for 13 pests of winter wheat, maize and oilseed rape and 17 pathogens of winter wheat, winter barley, oilseed rape, sugar beet and potatoes were analyzed (Tab. SI. 1).

### Landscape composition data

The French implementation of the registration for agricultural subsidies (RPG) in the framework of the European Common Agricultural Policy provides detailed annual information on the cropland cover over the French territory. The geometry of the fields is described by farmers based on the aerial photographs of the BD Ortho^®^. From 2006 to 2014 the description of the fields was by islet, a group of contiguous fields but 80% of them had only one type of crop. In each islet, the detailed surfaces were given by crop types (28 crop types for 329 crops registered). Here we use five of them: winter wheat, oilseed rape, winter barley, maize (including both silage and grain maize), other industrial crops (mainly and considered here to be beet) and flowering vegetables (mainly and considered here to be potatoes). From 2015 to 2017, both limitations have been lifted and the exact crop is known at the field level. We discarded any epidemiological observation whose recorded crop didn’t match the RPG.

The semi-natural elements considered were woods, grasslands, scrublands and hedgerows. The RPG provided us with grassland delineations the year of the observation (temporary and permanent are not distinguished), the BD TOPO^®^ (vegetation layer, version 2.1, 2017) provided us with the geometry of the other elements, considered stable in time. From this database, we group as “woodlands” the forests of broad-leaved, conifer, mixed species, with closed or open canopy.

### Variables calculations

The outbreak risk for each site.year.pest was defined as the rate of observations above the median of the observations for this pest for all site.year (Tab. SI 2). We quantified the landscape composition by measuring the surface (m^2^) of semi-natural elements and of the pest host crop around each observation in buffers of 200 m, 1 km, 5 km, and 10 km. To avoid attributing to the landscape the effects of the crop rotation at the plot scale, we take into account the latter through the time since the host crop on the one hand and grassland on the other hand have been grown on the plot. As only 2 years of RPG data were available before the first observation, we simplified this variable to three values: 1, 2 and 3 years or more. We discarded the points when the host crop or the grassland was not alone in the islet the last time it appeared. To account for the production environment, the farming type region was included as a factor (SI 4, Fig. SI 2a). To account for yearly weather we also included as factor the interaction between year and climatic zone^34^ (SI 4, Fig. SI 2b).

### Statistical analysis

For each pest, we described the outbreak risk with a generalized linear binomial model with a logit link^35^ as a function of the landscape, field level rotation and control factors. Models were fitted by LASSO penalized regressions using the glmnet R package^28^. The value of the penalization factor was set conservatively by cross-validation (10 folds) keeping the prediction error within one standard deviation of the minimum standard error (glmnet lambda1se). To limit the impact of random variations linked to the cross-validation procedure, the values considered for each pest in Figure 1 are medians of a 1000 bootstrap replicates (Fig. 1). When several scales were found significant for an element/pest pair, we only represent the most significant: the one with the largest effect size (the fits were realized on standardized parameter values). Binomial tests compared the repartition of negatively and positively correlated parameters for the different pests assuming a default binomial distribution with an equal probability of the two categories, two-sided p-values are used. To compare effect sizes odds ratios and risk ratios where assessed from the model estimated with the full dataset: landscape elements effect sizes were computed setting the selected scale at its 5th or 95th quantile among all observed year × site and setting all other parameters to their mean values. Year effect sizes were estimated similarly comparing for each pest and climatic region the best to the worst year and then taking the geometric mean over regions.

## Supporting information

All Supplementary materials

## Acknowledgments

We thank Marie-Hélène Jeuffroy for her help in designing the study; Danièle Simmoneau, Céline Gouwie, François Brun, Eric Cahuzac and Olivier Thérond for their support and expertise on the data; Felix Bianchi, Joséphine Python and Nathalie Cassagne for their feedback on early versions of the manuscript; Nicolas Guérin and Ryohei Chiyojima for participating in the analysis. We thank the IGN for providing the BD TOPO^®^ as part of a research agreement, the “agence de service et de paiement” for providing the RPG and the “conseil d’épidémiosurveillance” and Institut Technique de la Betterave for giving us access to epidemiological data. Funding were provided by the GIS GCHP2E and by the API-SMAL grant overseen by the French National Research Agency (ANR) as part of the “Investissements d’Avenir” Programme (LabEx BASC; ANR-11-LABX-0034).

## Notes

#### Summary of Updates

Much shorter version, formated as a letter, it also runs a bootstrap on the lasso analysis to assure the conclusions are stable across resampling of the initial dataset. The conclusions remain unchanged.

## References

1. Intergovernmental Science-Policy Platform on Biodiversity and Ecosystem Services, IPBES. The IPBES regional assessment report on biodiversity and ecosystem services for Europe and Central Asia. https://zenodo.org/record/3237429 (2018) doi:10.5281/zenodo.3237429.

2. DiBartolomeis, M., Kegley, S., Mineau, P., Radford, R. & Klein, K. An assessment of acute insecticide toxicity loading (AITL) of chemical pesticides used on agricultural land in the United States. PLOS ONE 14, e0220029 (2019).

3. Bass, C., Denholm, I., Williamson, M. S. & Nauen, R. The global status of insect resistance to neonicotinoid insecticides. Pestic. Biochem. Physiol. 121, 78–87 (2015).

4. Ferron, P. & Deguine, J.-P. Crop protection, biological control, habitat management and integrated farming. A review. Agron. Sustain. Dev. 25, 17–24 (2005).

5. Skellern, M. P. & Cook, S. M. Prospects for improved off-crop habitat management for pollen beetle control in oilseed rape. Arthropod-Plant Interact. 12, 849–866 (2018).

6. Yang, L. et al. Landscape structure alters the abundance and species composition of early-season aphid populations in wheat fields. Agric. Ecosyst. Environ. 269, 167–173 (2019).

7. Margosian, M. L., Garrett, K. A., Hutchinson, J. M. S. & With, K. A. Connectivity of the American Agricultural Landscape: Assessing the National Risk of Crop Pest and Disease Spread. BioScience 59, 141–151 (2009).

8. Woltz, J. M., Isaacs, R. & Landis, D. A. Landscape structure and habitat management differentially influence insect natural enemies in an agricultural landscape. Agric. Ecosyst. Environ. 152, 40–49 (2012).

9. Bianchi, F. J. J. A., Booij, C. J. H. & Tscharntke, T. Sustainable pest regulation in agricultural landscapes: a review on landscape composition, biodiversity and natural pest control. Proc. R. Soc. B Biol. Sci. 273, 1715–1727 (2006).

10. Karp, D. S. et al. Crop pests and predators exhibit inconsistent responses to surrounding landscape composition. Proc. Natl. Acad. Sci. 115, E7863–E7870 (2018).

11. Plantegenest, M., Le May, C. & Fabre, F. Landscape epidemiology of plant diseases. J. R. Soc. Interface 4, 963–972 (2007).

12. Sirami, C. et al. Increasing crop heterogeneity enhances multitrophic diversity across agricultural regions. Proc. Natl. Acad. Sci. 116, 16442–16447 (2019).

13. Kim, K.-H., Kabir, E. & Jahan, S. A. Exposure to pesticides and the associated human health effects. Sci. Total Environ. 575, 525–535 (2017).

14. Altieri, M., Nicholls, C. & Nicholls, C. Biodiversity and Pest Management in Agroecosystems. (CRC Press, 2018). doi:10.1201/9781482277937.

15. Chaplin-Kramer, R., O’Rourke, M. E., Blitzer, E. J. & Kremen, C. A meta-analysis of crop pest and natural enemy response to landscape complexity: Pest and natural enemy response to landscape complexity. Ecol. Lett. 14, 922–932 (2011).

16. Tscharntke, T., Klein, A. M., Kruess, A., Steffan-Dewenter, I. & Thies, C. Landscape perspectives on agricultural intensification and biodiversity – ecosystem service management. Ecol. Lett. 8, 857–874 (2005).

17. Veres, A., Petit, S., Conord, C. & Lavigne, C. Does landscape composition affect pest abundance and their control by natural enemies? A review. Agric. Ecosyst. Environ. 166, 110–117 (2013).

18. Perez-Alvarez, R., Nault, B. A. & Poveda, K. Contrasting effects of landscape composition on crop yield mediated by specialist herbivores. Ecol. Appl. 28, 842–853 (2018).

19. Tscharntke, T. et al. When natural habitat fails to enhance biological pest control – Five hypotheses. Biol. Conserv. 204, 449–458 (2016).

20. Yuen, J. & Mila, A. Landscape-Scale Disease Risk Quantification and Prediction. Annu. Rev. Phytopathol. 53, 471–484 (2015).

21. Aubertot, J.-N. & Robin, M.-H. Injury Profile SIMulator, a Qualitative Aggregative Modelling Framework to Predict Crop Injury Profile as a Function of Cropping Practices, and the Abiotic and Biotic Environment. I. Conceptual Bases. PLOS ONE 8, e73202 (2013).

22. Thies, C., Steffan-Dewenter, I. & Tscharntke, T. Interannual landscape changes influence plant– herbivore–parasitoid interactions. Agric. Ecosyst. Environ. 125, 266–268 (2008).

23. Eickermann, M., Beyer, M., Goergen, K., Hoffmann, L. & Junk, J. Shifted migration of the rape stem weevil Ceutorhynchus napi (Coleoptera: Curculionidae) linked to climate change. Eur. J. Entomol. 111, 243–250 (2014).

24. Jourdheuil, P. Remarques sur le nombre de générations de quelques Phyllotreta [Col. Chrysomelidae]. Bull. Société Entomol. Fr. 65, 126–131 (1960).

25. Ratnadass, A., Fernandes, P., Avelino, J. & Habib, R. Plant species diversity for sustainable management of crop pests and diseases in agroecosystems: a review. Agron. Sustain. Dev. 32, 273–303 (2012).

26. Rusch, A. et al. Agricultural landscape simplification reduces natural pest control: A quantitative synthesis. Agric. Ecosyst. Environ. 221, 198–204 (2016).

27. Lechenet, M., Dessaint, F., Py, G., Makowski, D. & Munier-Jolain, N. Reducing pesticide use while preserving crop productivity and profitability on arable farms. Nat. Plants 3, 17008 (2017).

28. Friedman, J., Hastie, T. & Tibshirani, R. Regularization Paths for Generalized Linear Models via Coordinate Descent. J. Stat. Softw. 33, (2010).

29. Chiverton, P. A. Predation of Rhopalosiphum padi (Homoptera: Aphididae) by polyphagous predatory arthropods during the aphids’ pre-peak period in spring barley. Ann. Appl. Biol. 111, 257–269 (1987).

30. Martin, E. A. et al. The interplay of landscape composition and configuration: new pathways to manage functional biodiversity and agroecosystem services across Europe. Ecol. Lett. (2019) doi:10.1111/ele.13265.

31. Lacasella, F. et al. From pest data to abundance-based risk maps combining eco-physiological knowledge, weather, and habitat variability. Ecol. Appl. 27, 575–588 (2017).

32. Schneider, G., Krauss, J., Riedinger, V., Holzschuh, A. & Steffan-Dewenter, I. Biological pest control and yields depend on spatial and temporal crop cover dynamics. J. Appl. Ecol. 52, 1283–1292 (2015).

33. Sine, M., Morin, E., Simonneau, D., Cosnac, G. D. & Escriou, H. VIGICULTURES – An early warning system for crop pest management. 8 (2010).

34. Lorgeou, J., Piraux, F., Picard, A. & Noël, V. Les grands contextes de production du blé tendre caractérisés par leurs stress. Perpective agricol. 22–24 (2012).

35. Guisan, A. & Zimmermann, N. E. Predictive habitat distribution models in ecology. Ecol. Model. 135, 147–186 (2000).

