## Supplementary material for "Higher surfaces of a crop in the landscape increase outbreak risks the following growing season": All Supplementary materials

### Supplementary information

#### *SI.1. Spatial distribution of the monitored fields*

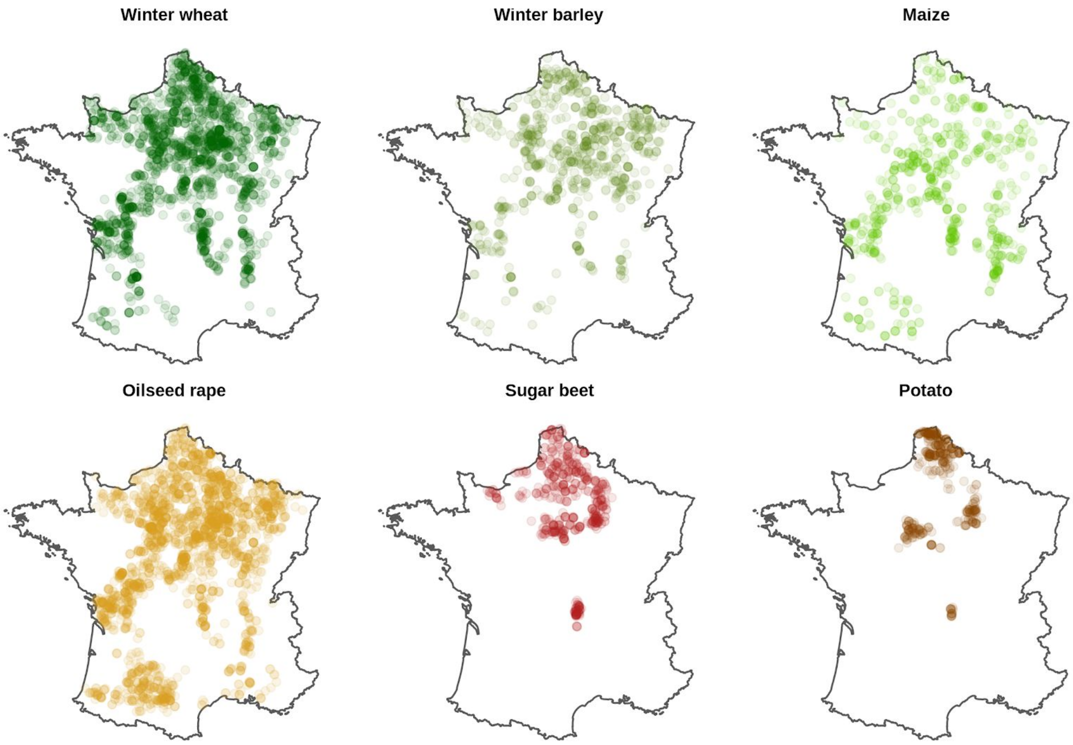


**Figure SI.1.** Spatial distribution of the monitored agricultural plots retained in the analysis for winter wheat (8 pathogens and 4 pests, 2246 field.year), winter barley (2 pathogens, 884 field.year), maize (1 pest, 745 field.year), oilseed rape (2 pathogens and 8 pests, 2617 field.year), sugar beet (4 pathogens, 572 field.year) and potato (1 pathogen, 411 field.year).

### *SI.2. Pests and pathogens monitoring details*

**Table SI. 1a**. Observation type, number and period for each of the animal pest studied

| Crop species | Bioagressor group[1] | Observation period[2] | N (plot × year) |
|---|---|---|---|
| Winter Wheat | *Sitodiplosis mosellan* | March-June | 631 |
| | *Deroceras, Arion, Limax spp.* | October-May | 1410 |
| | *Rhopalosiphum padi* | October-May | 1323 |
| | *Sitobion avenae* | March-August | 1154 |
| Maize | *Ostrinia nubilalis* | April-October | 745 |
| Oilseed rape | *Brevicoryne brassicae* | January-August | 2134 |
| | *Ceuthorhynchus napi* | January-May | 2343 |
| | *Ceutorhynchus assimilis* | February-August | 2162 |
| | *Ceutorhynchus picitarsis* | September-January | 2139 |
| | *Meligethes aeneus* | January-June | 2252 |
| | *Myzus persicae* | August-December | 1800 |
| | *Phyllotreta nemorum* | September-December | 1507 |
| | *Psylliodes chysocephala* | August-December | 2027 |

[1]Can be species, family or gender or of the bioagressor of interest
[2]Observations performed by Vigicultures ® experts during the 2009-2017 period

**Table SI. 1b**. Observation type, number and period for each of the pathogen studied

| Crop species | Bioagressor group[1] | Observation period[2] | N (plot × year) |
|---|---|---|---|
| Winter Wheat | *Blumeria graminis* | February-July | 1793 |
| | *Fusarium Graminearum* | February-July | 978 |
| | *Gaeumannomyces graminis* | March-July | 542 |
| | *Helminthosporium spp.* | February-July | 1160 |
| | *Oculimacula spp.* | February-July | 1411 |
| | *Puccinia striiformis* | January-July | 1622 |
| | *Puccinia triticina* | January-July | 1913 |
| | *Septoria tritici* | February-July | 2116 |
| Winter Barley | *Helminthosporium spp.* | February-July | 775 |
| | *Rhynchosporium secalis* | February-July | 790 |
| Oilseed rape | *Leptosphaeria maculans* | February-August | 1702 |
| | *Sclerotinia sclerotiorum* | March-June | 766 |
| Sugar Beet | *Cercospora beticola* | June-October | 572 |
| | *Erysiphe betae* | June-October | 567 |
| | *Ramularia betae* | June-October | 568 |
| | *Uromyces betae* | June-October | 567 |
| Potatoes | *Phytophthora infestans* | April-September | 411 |

[1]Can be species, family or gender or of the bioagressor of interest
[2]Observations performed by Vigicultures ® experts during the 2009-2017 period

### *SI.3. Pests and pathogens observation metrics and outbreak threshold*

**Table SI. 2a**. Observation type, number and period for each of the pest studied

| Crop species | Bioagressor group[1] | Observation metric | Thr. low. | Thr mid. | Thr. high |
|---|---|---|---|---|---|
| Winter Wheat | *Sitodiplosis mosellan* | # [2] observed in yellow bowl | 0.00 | 0.67 | 2.00 |
| | *Deroceras, Arion, Limax spp.* | % of seedlings with damages | 0.00 | 0.00 | 0.00 |
| | *Rhopalosiphum padi* | % of plants with insect present | 0.00 | 0.00 | 0.00 |
| | *Sitobion avenae* | % of plants with insect present | 0.00 | 0.00 | 0.00 |
| Maize | *Ostrinia nubilalis* | # adults in pheromone traps | 0.00 | 1.78 | 6.00 |
| Oilseed rape | *Brevicoryne brassicae* | # colony per $m^2$ | 0.00 | 0.00 | 0.00 |
| | *Ceuthorhynchus napi* | # captured in traps | 0.00 | 1.60 | 5.00 |
| | *Ceutorhynchus assimilis* | # per plants | 0.00 | 0.01 | 0.05 |
| | *Ceutorhynchus picitarsis* | # captured in traps | 0.00 | 0.57 | 2.00 |
| | *Meligethes aeneus* | % of plants with insect present | 0.00 | 20.00 | 50.00 |
| | *Myzus persicae* | % of plants with insect present | 0.00 | 0.00 | 0.00 |
| | *Phyllotreta nemorum* | # captured in vegetation traps | 0.00 | 0.00 | 0.00 |
| | *Psylliodes chysocephala* | # captured in ground traps | 0.00 | 1.50 | 4.00 |

[1]Can be species, family or gender or of the bioagressor of interest
[2]# : number counted

**Table SI. 2b**. Observation type, number and period for each of the pathogen studied

| Crop species | Bioagressor group[1] | Observation metric | Thr. low | Thr. mid. | Thr. high |
|---|---|---|---|---|---|
| Winter Wheat | *Blumeria graminis* | Severity scale 1:10 [2] | 0.00 | 0.00 | 0.00 |
| | *Fusarium Graminearum* | % of the base stem infected | 0.00 | 0.00 | 0.00 |
| | *Gaeumannomyces graminis* | Severity scale 1:100 | 0.00 | 0.00 | 0.00 |
| | *Helminthosporium spp.* | Severity scale 1:10 [2] | 0.00 | 0.00 | 0.00 |
| | *Oculimacula spp.* | Severity scale 1:100 | 0.00. | 0.00 | 0.00 |
| | *Puccinia striiformis* | Severity scale 1:100 [2] | 0.00 | 0.00 | 0.00 |
| | *Puccinia triticina* | Severity scale 1:10 [2] | 0.00 | 0.00 | 0.00 |
| | *Septoria tritici* | Severity scale 1:10 [2] | 0.00 | 2.31 | 6.00 |
| Winter Barley | *Helminthosporium spp.* | Severity scale 1:10 [2] | 0.00 | 1.50 | 3.00 |
| | *Rhynchosporium secalis* | Severity scale 1:10 [2] | 0.00 | 1.17 | 3.00 |
| Oilseed rape | *Leptosphaeria maculans* | % of plants with stem necrosis | 0.00 | 0.00 | 0.00 |
| | *Sclerotinia sclerotiorum* | % of flowers affected | 45.00 | 45.00 | 47.50 |
| Sugar Beet | *Cercospora beticola* | % of infected leaves | 0.00 | 2.79 | 9.00 |
| | *Erysiphe betae* | % of infected leaves | 0.00 | 0.00 | 0.00 |
| | *Ramularia betae* | % of infected leaves | 0.00 | 0.25 | 1.00 |
| | *Uromyces betae* | % of infected leaves | 0.00 | 2.18 | 8.00 |
| Potatoes | *Phytophthora infestans* | Severity scale 1:10 | 0.00 | 0.00 | 0.00 |

[1]Can be species, family or gender or of the bioagressor of interest
[2]On the third leaf (F3)

*SI.4 Definition of the regions for the control variables*

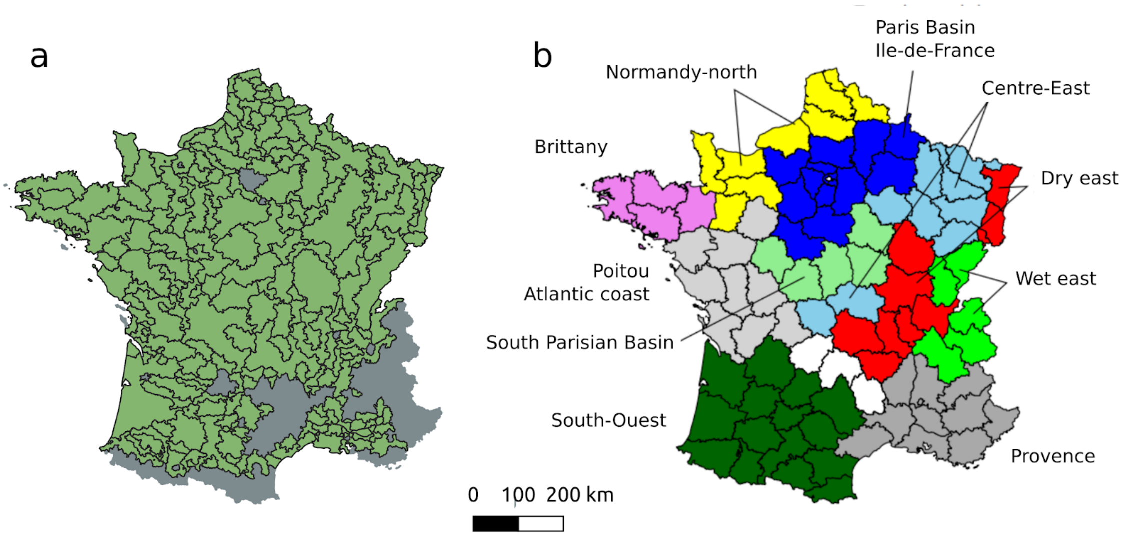


**Figure SI.2** Control regions for farming system region (a) and agro-climatic regions (b).
In (a), the farming system regions are contiguous green regions; grey areas are urban or mountainous regions where few to no annual crops are found. In (b), the agro-climatic regions[1] can be discontinuous but have the same color. They are defined by aggregating administrative departments whose boundaries are superimposed.

[1] Lorgeou, J., Piraux, F., Picard, A., ARVALIS - Institut du Végétal, et Noël, V., 2012. Les grands contextes de production du blé tendre caractérisés par leurs stress. Perpective agricole, n° 389. p. 22-24

*SI.5 Sensitivity analysis to alternative formulations*

To evaluate the robustness of our results, we tested alternative models modifying either their mathematical expression or their input variables (Fig. SI 3). In the reference model we systematically used a log transform of the input variables. We also tested a model without this log transform. Overall, this induced little change to the tendencies.

A possible assumption in biological control by conservation is that it would avoid the most outrageous outbreaks. This implies that using higher thresholds of abundance in our models should increase the positive impact of semi-natural elements. However, the few differences between the low, intermediate and high threshold suggest here no such relationship though we only differentiate such thresholds for a minority of pests and diseases (Table SI 2a,b).

Despite possible correlation between the abundance of a crop in the rotation and its abundance in the landscape, removing the rotation on the field from the available variables didn't affect in the least the correlations with the crops in the landscape, confirming those variables are largely independent in our reference model. This could be nevertheless different if we had not the farming system region as a control variable. Indeed, the model without it sees a reinforcement of the importance of the in-field rotation for pathogens and some modifications of the tendencies in the correlations with host crop surfaces the current and past growing seasons.

As the landscape at 10km could also be correlated with broad agronomic conditions changes, we assessed the robustness of the conclusions derived from the model when accounting for the PCA components of the 10km landscape elements as a continuous version of the farming system environment and removing the original input variable associated with the 10km buffer variables. Logically, this removed the correlation with the host crop the previous year for the three pathogens with a maximum correlation at 5 or 10km but hardly changed the patterns otherwise. Finally, the control for regional weather conditions that massively morcellate the analysis may be either decreasing the statistical power if not justified or increase it if it allows to account for important corrections removing a lot of meteorological condition noise from the data. Eight of the eleven tested elements see a decrease of the detected correlations when we remove this control, suggesting that accounting for weather variability tend to increase the ability to detect the impact of landscape elements.

Note that in the Figure SI.3 models are fit on the whole dataset, not using the bootstrap procedure explaining slight differences between the first line (Reference) and the results presented in Figure 1 which represent the best estimate over all the bootstrapped samples (see SI 7 to apprehend the variability in the bootstrapped samples).

a)

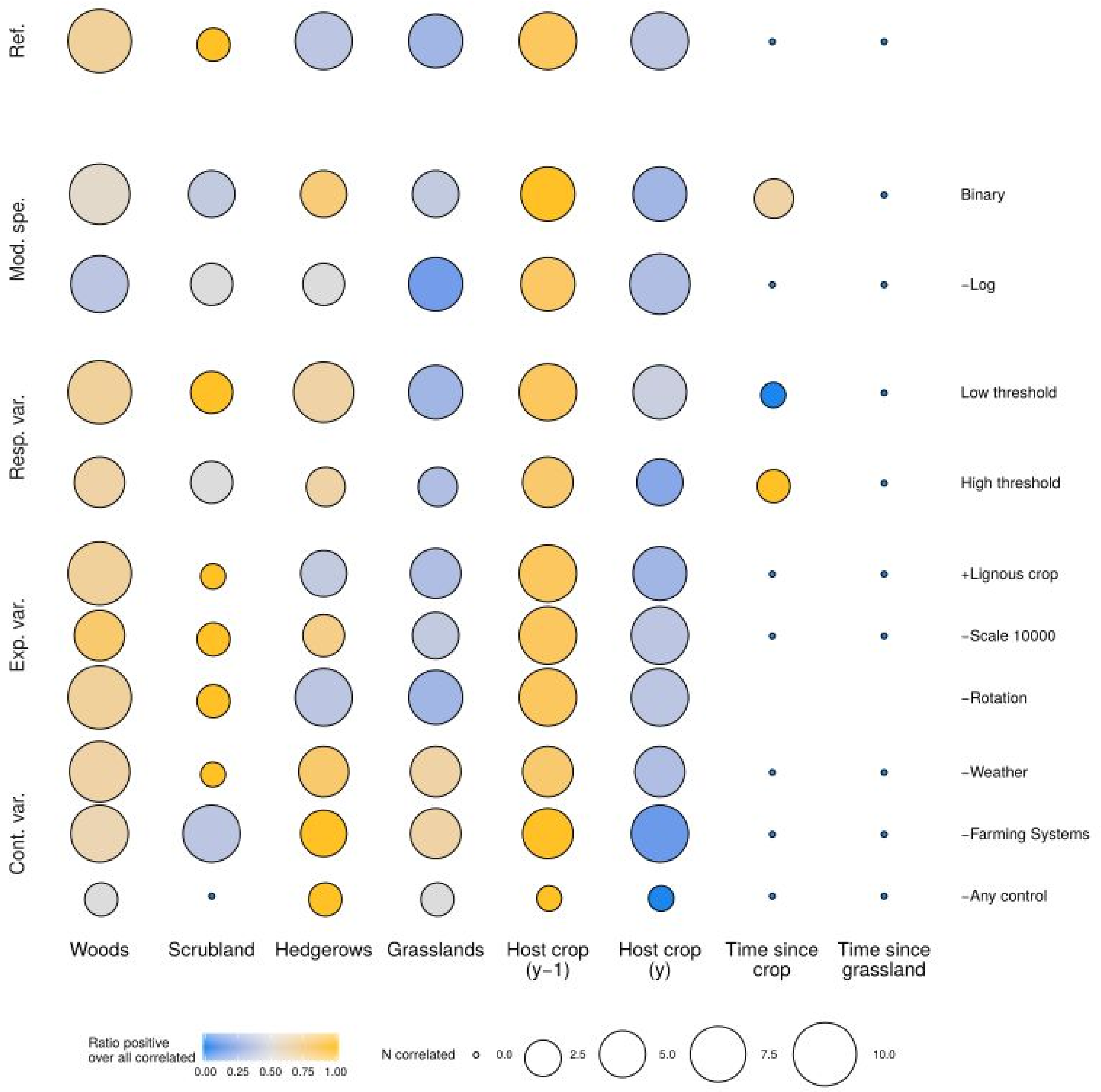


**Figure SI.3**. Summary of the landscape components and crop rotation effect on a) animal pests b) pathogens and c) all organisms, according to model variations. The color indicates the ratio of organisms with positive correlation over all the organisms with positive correlation. N correlated: number of pests positively or negatively correlated. The correlation used for each organism and landscape element is the scale correlation with the largest absolute value. *Ref.*: full model used in the main text. *Mod. spe.*: changing model specification i.e. *Binary*: presence-absence over the year; *-Log*: do not take the log of the parameter data before fitting. *Resp. var.*: response variable i.e., *Low/high threshold*: changing outbreak threshold (Tab. SI2). *Exp. var.*: changing explaining variables i.e. *+Lignous crop*: adding vineyards and orchards as defined in the BD TOPO vegetation layer as landscape determinants; *-Scale 10000*: replacement of the 10 km scale for all elements by the 4 principal components on this scale for all elements; *-Rotation*: removing the Time since crop and Time since grassland component. *Cont. var.*: changing control variables i.e. *-Weather*: removing the interaction between climatic region and year (no year or climatic region factor left either); -*Farming Systems*: removing the farming system regions control variable; -*Any control*: removing both weather and farming system regions.

b)

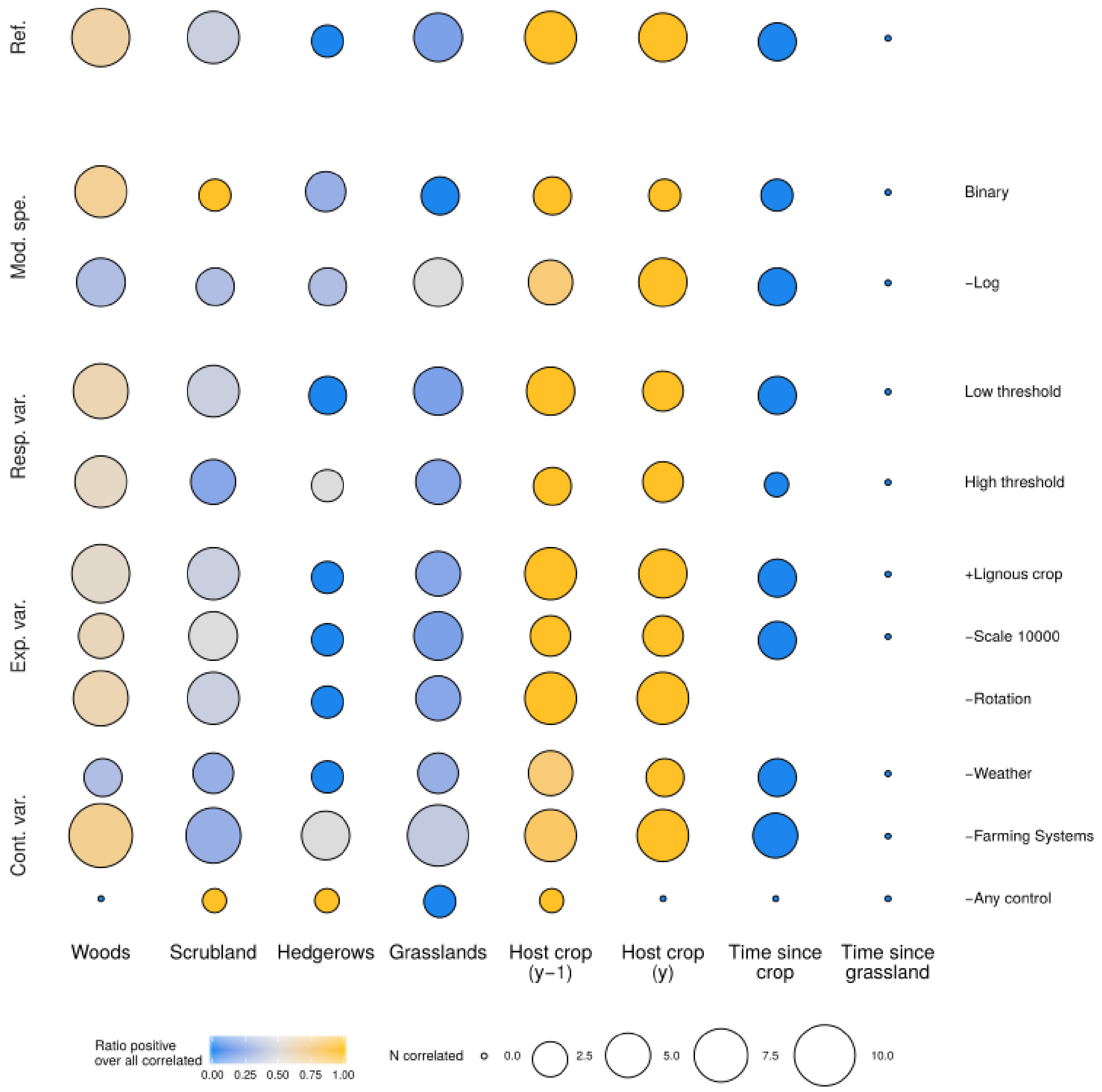

c)

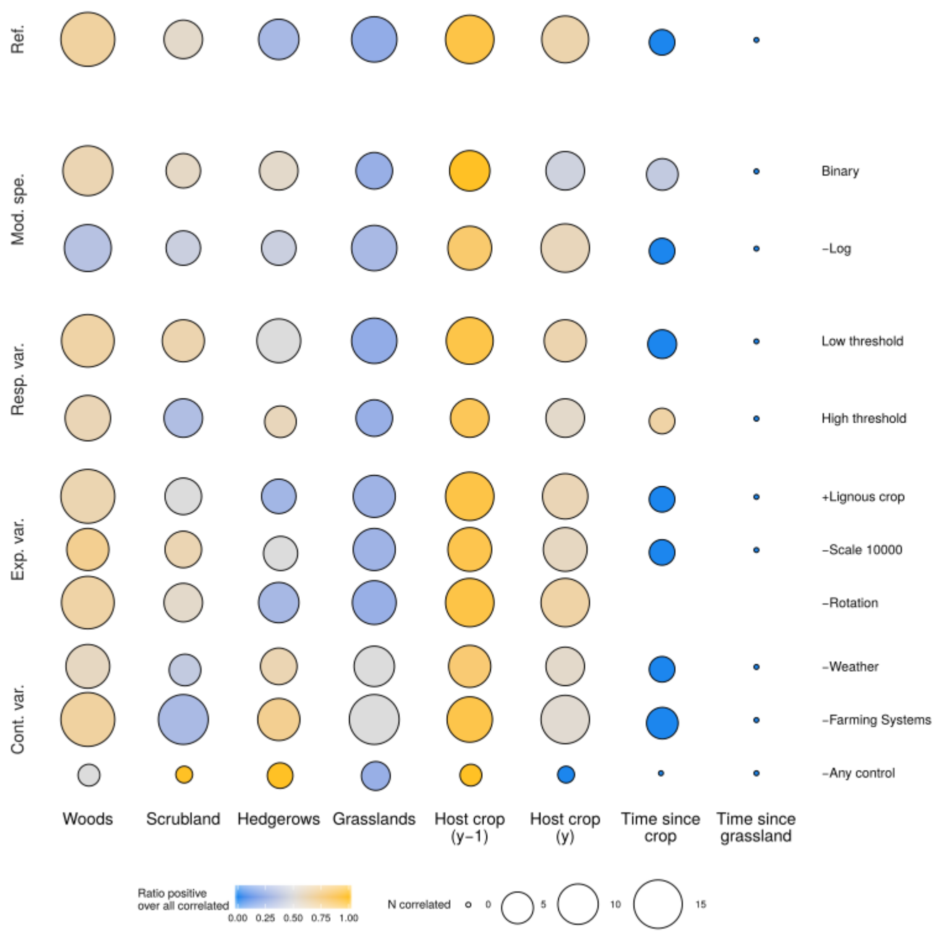
Ref.
Mod. spe.
Resp. var.
Exp. var.
Cont. var.
Binary
-Log
Low threshold
High threshold
+Lignous crop
-Scale 10000
-Rotation
-Weather
-Farming Systems
-Any control
Woods
Scrubland
Hedgerows
Grasslands
Host crop (y-1)
Host crop (y)
Time since crop
Time since grassland
Ratio positive over all correlated
0.00 0.25 0.50 0.75 1.00
N correlated
0 5 10 15

*SI.6:* **Figure SI.4, estimates by organism**

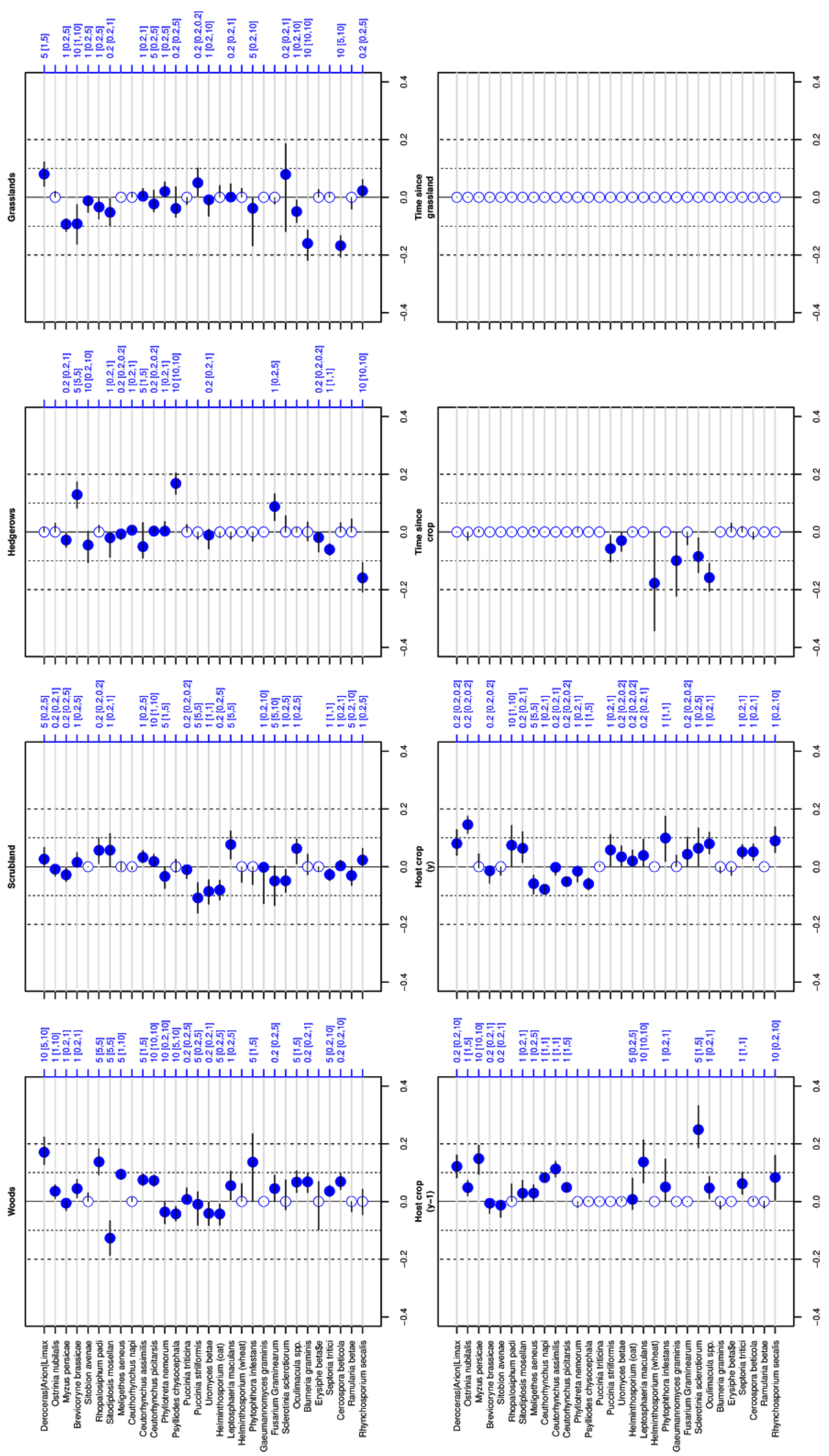


Blue point : median over a 1000 bootstrap samples of the largest coefficient estimate (as absolute value) over the four possible scales by pest. Variables are normalized before the fit to allow the comparison between the coefficients. The segments indicate the corresponding 25% and 75% quantiles. The right blue axis corresponds to the median selected distance as well as the 25%-75% quantile. The pests are ranked by type (animal, pathogens) and order. Vertical dashed lines only mark values 0.1 and 0.2.

*SI.7 Observed distribution in the bootstrap procedure*

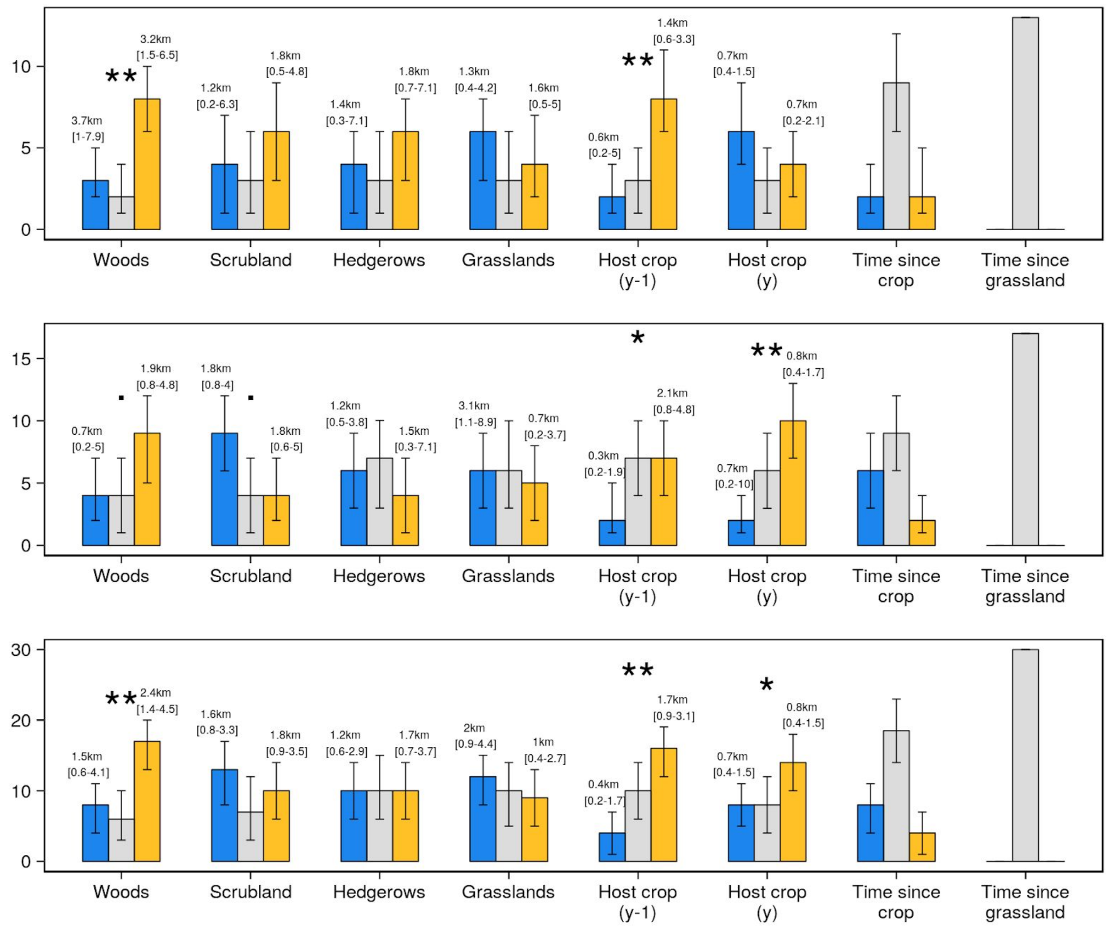


**Figure SI.5** : Distribution of the results in a thousand bootstrapped samples. a, animal pests; b, pathogens; c, total. Blue negative coefficients, grey null coefficients, orange positive coefficients, the bars indicate the median values, the moustache indicate the 2.5 and 97.5% quantiles. The stars correspond to the probability that the direction of the difference between positive and negative coefficients is lost in our 1000 bootstrap samples. .:p.<0.1; *: p. <0.05; **: p. <0.01.

Note that while the Figure 1 displays a summary of the best estimate (median) for each pest, this figure displays the distribution of the number of organisms with positive, negative and zero coefficients over the 1000 bootstrap samples. Consequently, it answers a different question: in the figure 1, a significant test indicates, given our best guess for each pest, a tendency likely to be observed over the different pests of the same group than the pests observed while in this figure, here, a significant test indicate that the tendency is well supported for the observed pests by our data. In particular, while the observed difference between positive and negative correlations with the wood is very strongly supported by our data (Figure SI.5), the number of organisms positively vs. negatively correlated is not enough to strongly suggest a general tendency over the animal pests or all pests (Figure 1).

Note: in our bootstrap procedure, the occurrence of samples not leading to a ratio of positive/(negative+positive) higher for pathogens than for animal pests was less than 0.5%.
